# Homologous recombination induced by a replication fork barrier requires cooperation between strand invasion and strand annealing activities

**DOI:** 10.1101/2021.08.20.456128

**Authors:** Léa Marie, Lorraine S. Symington

## Abstract

Replication stress and abundant repetitive sequences have emerged as primary conditions underlying genomic instability in eukaryotes. Elucidating the mechanism of recombination between repeated sequences in the context of replication stress is essential to understanding how genome rearrangements occur. To gain insight into this process, we used a prokaryotic Tus/*Ter* barrier designed to induce transient replication fork stalling near inverted repeats in the budding yeast genome. Remarkably, we show that the replication fork block stimulates a unique recombination pathway dependent on Rad51 strand invasion and Rad52-Rad59 strand annealing activities, as well as Mph1/Rad5 fork remodelers, Mre11/Exo1 short and long-range resection machineries, Rad1-Rad10 nuclease and DNA polymerase δ. Furthermore, we show recombination at stalled replication forks is limited by the Srs2 helicase and Mus81-Mms4/Yen1 structure-selective nucleases. Physical analysis of replication-associated recombinants revealed that half are associated with an inversion of sequence between the repeats. Based on our extensive genetic characterization, we propose a model for recombination of closely linked repeats at stalled replication forks that can actively contribute to genomic rearrangements.

## INTRODUCTION

Maintaining genome integrity is essential for accurate transmission of genetic information and cell survival. Replication stress has emerged as a major driver of genomic instability in normal and cancer cells. Replication forks become stressed as a result of DNA lesions, spontaneous formation of secondary structures, RNA-DNA hybrids, protein-DNA complexes, activation of oncogenes, or depletion of nucleotides ^1–3^. These obstacles to the progression of replication can cause forks to slow down, stall and collapse. Consequently, multiple mechanisms have evolved to handle perturbed replication forks to ensure genomic stability ^4^.

In eukaryotes, the presence of multiple replication origins, including dormant origins that are fired in response to replication stress, is one way to ensure complete genome duplication ^5,6^. Alternatively, the obstacle can be bypassed by translesion polymerases or by legitimate template switching. The latter is a strand exchange reaction mediated by homologous recombination (HR) proteins, consisting of annealing a nascent strand to its undamaged sister chromatid to template new DNA synthesis ^7^. In recent years, replication fork reversal has also emerged as a central remodeling process in the recovery of replication in both eukaryotes and bacteria ^8–12^. This process allows stalled replication forks to reverse their progression through the unwinding and annealing of the two nascent strands concomitant with reannealing of the parental duplex DNA, resulting in the formation of a four-way-junction, sometimes called a chicken-foot structure. Consequently, the lesion can be bypassed by extension of the leading strand using the lagging strand as a template followed by branch migration of the reversed structure. Alternatively, the extruded nascent strands can undergo HR-dependent invasion of the homologous sequence in the reformed parental dsDNA, resulting in the formation of a D-loop to restart replication. In bacteria, the replisome is reassembled on the D-loop structure ^13^, whereas in eukaryotes DNA synthesis within the D-loop can extend to the telomere or be terminated by a converging replication fork ^5^. In addition, relocation of a lesion back into the parental duplex could facilitate repair by the excision repair pathways ^14^.

Thus, along with its critical role in DNA repair and segregation of chromosome homologs during meiosis, HR is involved in multiple replication restart mechanisms, which contribute to the preservation of genome integrity. However, HR can also be a source of instability as it occasionally occurs between chromosome homologs in diploid mitotic cells, resulting in loss of heterozygosity. Moreover, non-allelic HR (NAHR) between dispersed repeats can cause genome rearrangements ^15–18^. A significant factor underlying chromosome rearrangements is the abundance of repeated sequences in eukaryotic genomes. Approximately 45% of the human genome is composed of repetitive sequences including transposon-derived repeats, processed pseudogenes, simple sequence repeats, tandemly repeated sequences and low-copy repeats (LCRs) distributed across all chromosomes ^19,20^. NAHR between repeated sequences can lead to deletions, duplications, inversions or translocations ^21–27^. Consequently, NAHR has been associated with many genomic disorders ^28,29^ and is a major contributor to copy-number variation (CNV) in humans.

It is well established that rearrangements due to NAHR can result from the repair of double strand breaks (DSBs) ^30–34^. However, studies in yeast, human and bacteria have shown that such genomic alterations can also arise during replication ^18,24,35,36^. Notably, studies in *Schizosaccharomyces pombe* have shown that a protein-induced, site-specific replication fork barrier can cause a high frequency of genomic rearrangements in the absence of a long-lived DSB intermediate ^24,37^, consistent with the idea that replication stress contributes to NAHR. Elucidating the molecular mechanisms of NAHR occurring during the processing and restart of stressed replication forks remains crucial to understanding how genome rearrangements occur.

In *Saccharomyces cerevisiae*, spontaneous HR between repeated sequences shows different genetic requirements depending on the genomic location of the repeats. Inter-chromosomal recombination is generally Rad51 dependent, whereas recombination between tandem direct repeats can occur by Rad51-independent single-strand annealing (SSA) ^38^. It has been shown that repeats in inverted orientation can spontaneously recombine by Rad51-dependent and Rad51-independent mechanisms ^39^, and these two pathways generate different recombination products. Rad51-mediated recombination results in gene conversion, which maintains the intervening sequence in the original configuration, whereas Rad51-independent recombination leads to inversion of the intervening DNA. The inversion events require Rad52 and Rad59 ^40^, which are known to catalyze annealing of RPA-coated single-stranded DNA (ssDNA) *in vitro*, and are required for SSA *in vivo*. Because DSB-induced recombination between inverted repeats is dependent on Rad51 ^41^, it was proposed that the spontaneous Rad51-independent inversions could be the result of annealing between exposed ssDNA at stressed replication forks ^42^.

To elucidate the mechanism of NAHR between inverted repeats in the context of replication stress, we investigated the role of a protein-induced replication fork barrier in promoting inverted repeats recombination. Previous studies have shown that the *Escherichia coli* Tus/*Ter* complex can function as a DNA replication fork barrier when engineered into the genome of yeast or mouse cells ^43–45^. Here, we demonstrate that a polar replication fork barrier engineered to induce fork stalling downstream of inverted repeats is sufficient to trigger NAHR. Physical analysis of the recombinants showed that gene conversion and inversion events were stimulated to the same extent. Unlike spontaneous events, we found that replication-associated NAHR unexpectedly relies on a unique pathway dependent on Rad51 strand invasion and Rad52-Rad59 strand annealing activities. We discuss a model to account for dependence on both Rad51 and Rad52-Rad59 and formation of gene conversion or inversion outcomes.

## RESULTS

### A polar replication fork barrier stimulates NAHR

To assess NAHR, we used a recombination reporter composed of two *ade2* heteroalleles oriented as inverted repeats ^39^. The inverted repeat cassette was inserted at the *HIS2* locus, 4 kb centromere distal to the efficient *ARS607* replication origin, on chromosome 6. The origin-proximal *ade2-n* allele contains a +2 frameshift located 370 bp away from the stop codon and is transcribed by the native *ADE2* promoter. The origin-distal allele, *ade2Δ5’*, has a deletion of the first 176 nucleotides along with the promoter. The two repeats share 1.8 kb of homology and are separated by 1.4 kb containing a *TRP1* gene transcribed by its native promoter (Fig 1A).

**Figure 1.**
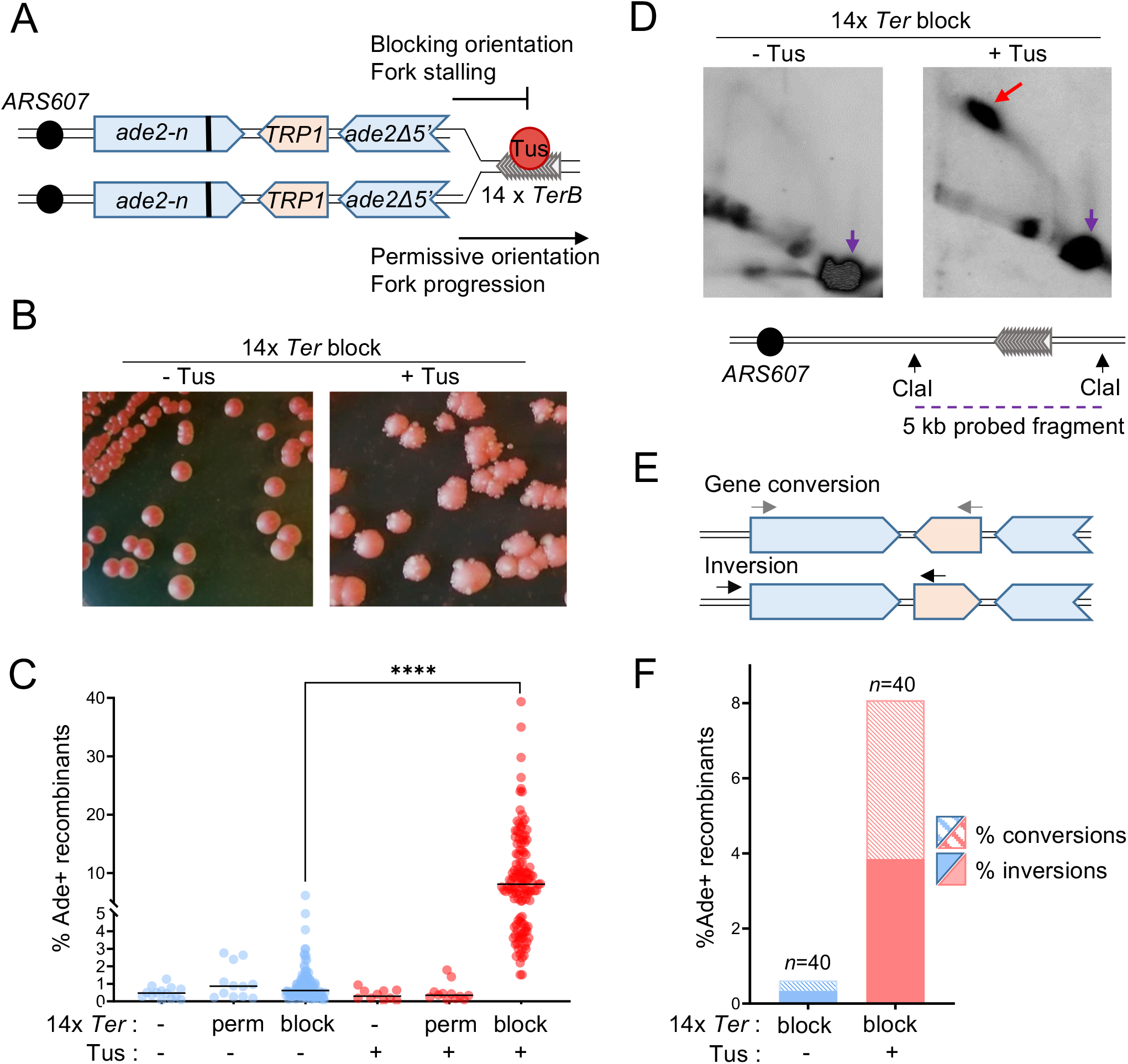
A localised fork stalling barrier stimulates NAHR. **A** Schematic of the *ade2* reporter and Tus/*Ter* barrier in the blocking or permissive orientation with regard to *ARS607*. The bold line indicates the +2 frame-shift mutation. **B** Colonies form more white sectors and papillae (indicative of Ade^+^ phenotype) when Tus expression is induced. block=blocking orientation. **C** Quantifications of Ade^+^ recombinants without (blue data points) and with (red data points) induction of Tus expression in strains containing different *Ter* constructs (see Table S1). Black lines indicate medians. P-values are reported as stars when significant: **** p-value <0.0001. Perm=permissive orientation; block=blocking orientation. **D** Two-dimensional gel analysis of replication intermediates in the strain containing 14 *Ter* repeats in the blocking orientation, with or without induction of Tus expression (see Fig S1 for details of probes used). The red arrow indicates fork arrest along the Y-shaped replication arc, the purple arrow indicates the unreplicated fragment. **E** Ade^+^ recombinants formed by gene conversion or by inversion of the *TRP1* locus are distinguished by PCR using primers designated by gray or black arrows. Inversion events can have the wild type or +2 frameshift site within the *ade2Δ5’* allele and are not distinguished here. **F** Distribution of NAHR events for each condition. *n* indicates the number of independent Ade^+^ recombinants tested.

To analyze recombination in the context of a unique stressed replication fork, in the absence of any genome-wide stress or global checkpoint activation, we took advantage of the galactose-inducible Tus/*Ter* replication fork barrier ^43,46^. We inserted 14 *TerB* repeats (hereafter referred to as 14 *Ter*) in the permissive or blocking orientation relative to *ARS607*, 120 bp or 170 bp distal to the *ade2Δ5’* repeat, respectively (Fig 1A). The location was selected based on a previous study showing that Tus/*Ter* induces mutagenesis of the newly replicated region behind the stalled fork ^47^. The *P_GAL1_-Tus* cassette was integrated at the *LEU2* locus.

In cells containing 14 *Ter* repeats in the blocking orientation, an elevated proportion of colonies developing white sectors, indicative of an Ade^+^ phenotype, was noticeable on plates containing galactose (Fig 1B). Consistently, quantification of Ade^+^ recombinants arising in this strain showed that expression of the Tus protein stimulated recombination frequency from 0.62% to 8.08% (Fig 1C; Table S1). We confirmed that the induction of the Tus protein expression had no effect on recombination frequency in cells containing no Ter repeats or 14 *Ter* repeats in the permissive orientation (Fig 1C; Table S1). By two-dimensional (2D) gel analysis of a 5 kb fragment encompassing part of the *ade2* reporter and the *Ter* repeats, we confirmed that induction of the Tus protein expression generates a significant replication fork arrest in the strain containing 14 *Ter* repeats in the blocking orientation (Fig 1D; Fig S1). Thus, replication fork stalling at a polar Tus/*Ter* barrier stimulates recombination between inverted repeats, more than 10-fold. We investigated the nature of the Tus/*Ter*-induced events by a PCR-based method (Fig 1E). Gene conversions and inversions were equivalently induced upon expression of the Tus protein, representing 47.5% and 52.5% of the Ade^+^ recombinants, respectively (Fig 1F).

The role of genome-wide replication stress in stimulation of NAHR was assessed by growing cells with the *ade2* reporter on media containing DNA damaging agents known to induce replication stress, namely, methyl methanesulfanate (MMS), camptothecin (CPT) or hydroxyurea (HU). Within three days, an increased proportion of colonies containing white sectors, indicative of an Ade^+^ phenotype, was clearly visible in the presence of MMS and CPT (Fig S2A). Consistently, quantification of Ade^+^ recombination frequencies under normal conditions (0.62% spontaneous recombination) and genotoxic conditions (16.15% with MMS, 9.79% with CPT, 1.5% with HU) revealed a strong stimulation of recombination between the inverted *ade2* repeats in presence of MMS and CPT (Fig S2B). The types of recombination events induced by MMS or CPT were determined by PCR analysis of independent recombinants. In the presence of MMS, the frequency of gene conversions was 30-fold higher (10.8%), whereas the frequency of inversions was increased by a factor 18 (5.4%). In presence of CPT, the frequency of gene conversions was 9 times higher (3.2%), whereas inversions were induced 22-fold (6.36%) (Fig S2C). We note that in the presence of CPT, the nature of the recombination event of a small proportion of recombinants could not be easily determined by the PCR method employed and these were not analyzed further. We detected a moderate induction of recombination frequency by HU and the distribution of gene conversions and inversions appeared similar to normal conditions (Fig S2B and C). Since replication fork arrest by a protein block is effective in stimulating NAHR, we suggest that the attenuated induction of Ade^+^ recombinants in response to HU is due to the dNTP requirement for DNA synthesis associated with recombination-dependent fork restart.

Together, these results indicate that NAHR between long inverted repeats, leading to gene conversion or inversion of the intervening sequence, can be generated by genome-wide replication stress or by a localized replication fork barrier, consistent with prior studies in *S. pombe* and mouse cells _24,37,44_.

### NAHR associated with replication fork stalling has unique genetic requirements compared to spontaneous NAHR

In line with previous studies ^40,42^, we found that spontaneous gene conversions and inversions are products of two independent recombination pathways. In the absence of Tus/*Ter*-induced replication stress, deletion of *RAD51* or *RAD59* only partially decreased recombination between the *ade2* repeats, whereas no recombination was detected in the double mutant. The Rad52 protein is involved in both pathways as the recombination frequency of the *rad52Δ* strain, like the *rad51Δ rad59Δ* double mutant, was below detection (Fig 2A, Table S1). Physical analysis of spontaneous Ade^+^ recombinants arising in the *rad51Δ* mutant showed that 82% of the tested recombinants contained an inversion. On the other hand, in the *rad59Δ* mutant, 71% of the recombinants were gene conversions (Fig 2B). These results confirm that spontaneous NAHR events leading to inversions of the *TRP1* gene are largely independent of Rad51 and require Rad59 and Rad52, whereas gene conversions are mostly independent of Rad59 and require Rad51 and Rad52 ^40^.

**Figure 2.**
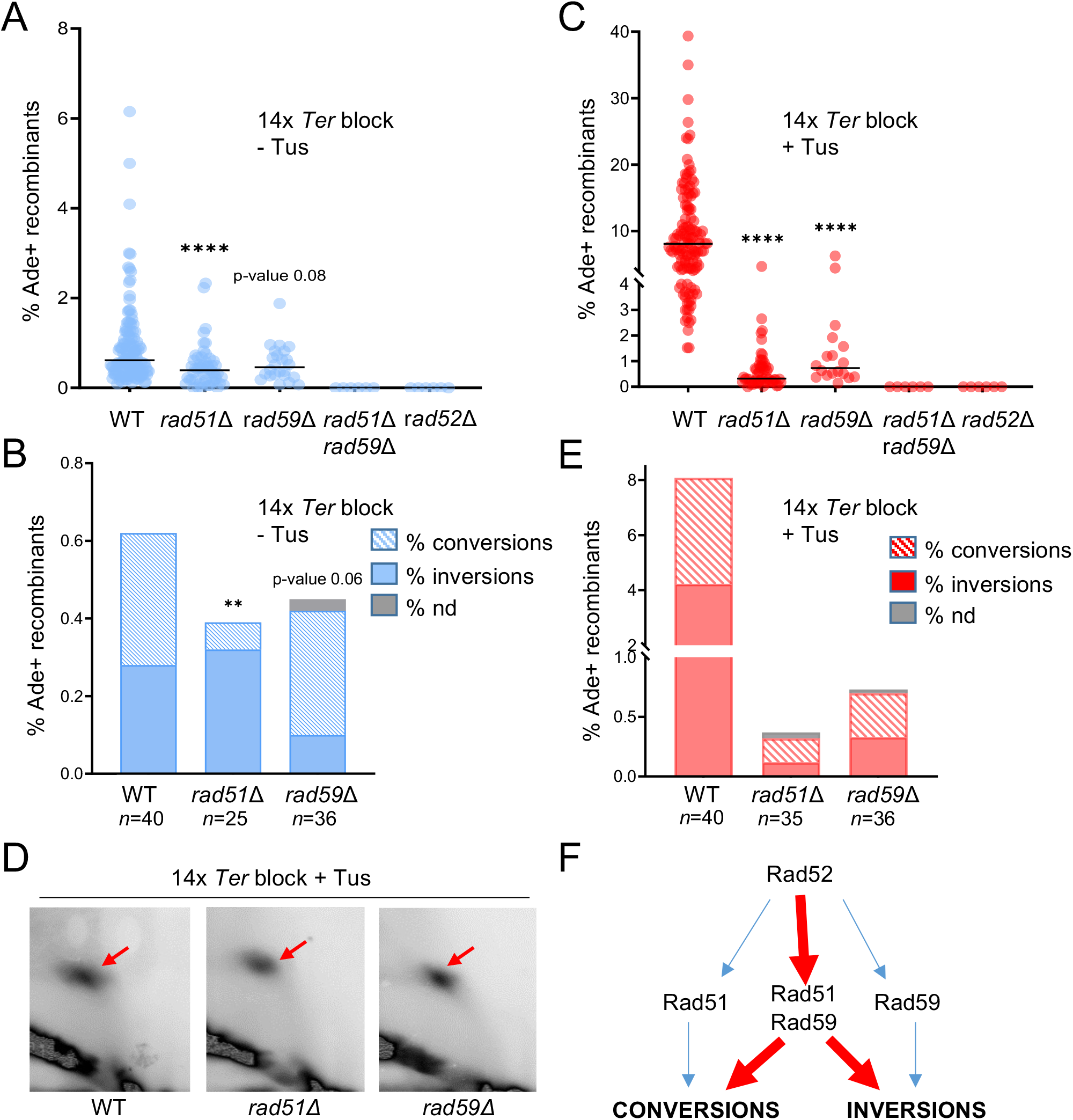
NAHR at the Tus/*Ter* barrier relies on the cooperation of Rad51 and Rad59. **A** Spontaneous Ade^+^ recombination frequencies in WT and mutant strains. Black lines indicate medians. P-values, reported as stars when significant, are relative to the WT strain in the same condition: **** p-value <0.0001. **B** Distribution of independent spontaneous recombination events scored by PCR; nd indicates structure could not be not determined by PCR; *n* indicates the number of independent Ade^+^ recombinants tested. P-values, reported as stars when significant, are relative to the WT strain: ** p-value <0.005 **C** Tus-induced Ade^+^ recombination frequencies in WT and mutant strains. **D** 2D gel analysis of replication intermediates showing similar fork arrest in the WT and mutant strains. Red arrows indicate the replication fork arrest on the arc of Y-shaped replication intermediates. **E** Distribution of events scored by PCR for Tus-induced recombinants. **F** Contribution of Rad51, Rad52 and Rad59 to spontaneous (blue) and replication fork block induced (red) NAHR pathways.

Surprisingly, both *rad51Δ* and *rad59Δ* single mutants showed no induction of recombination by the Tus/*Ter* replication barrier (Fig 2C). We confirmed by 2D gels that the Tus-generated replication fork barrier was still detected in both mutants (Fig 2D). Thus, it appears that gene conversions and inversions associated with replication fork stalling have specific genetic requirements. Rad52 is essential for recombination in this context as well since no Ade^+^ recombinants were detected in the *rad52Δ* strain (Fig 2C). Intriguingly, physical analysis of Tus-induced recombinants in the *rad51Δ* and *rad59Δ* mutant strains revealed a different distribution from spontaneous events (Fig 2E).

We also determined the frequency of MMS and CPT-stimulated recombination in *rad51Δ* and *rad59Δ* single mutants using low concentrations of the drugs that allowed growth of the mutants while stimulating recombination in the WT strain (Fig S2D). Whereas the frequency of recombination in the WT strain increased from 0.62% to 14.38% with MMS and 2.09% with CPT, we detected no stimulation of recombination by genotoxic agents in *rad51Δ* and *rad59Δ* mutants.

Together, these results indicate that recombination between inverted repeats associated with replication stress is mediated by a unique molecular mechanism involving Rad51, Rad52 and Rad59, and leads to both gene conversions and inversions (Fig 2F).

### Replication associated NAHR events rely on Rad51 strand invasion and Rad52 strand annealing activities

We next wanted to further explore the roles of Rad51 and Rad52 in Tus/*Ter*-induced NAHR. Rad51 has three established functions at stalled replication forks. First, Rad51 promotes replication fork reversal in mammalian cells, but does not have fork remodeling activity on its own and different models have been proposed to explain its role in this process ^8^. Second, Rad51 is required for protection of nascent DNA strands at reversed forks from extensive nucleolytic degradation by Mre11 ^48,49^. Finally, Rad51 plays a role in the restart of arrested replication forks by several recombination pathways involving strand invasion and strand exchange ^8,49,50^.

The *rad51-*II3A allele contains three amino acid substitutions, eliminating the secondary DNA binding site. The mutant protein retains the ability to form filaments on ssDNA but is defective for strand exchange activity ^51,52^. A recent study, modeling this mutation in human cells, revealed that the enzymatic activity of Rad51 is neither required to promote fork reversal nor to protect stalled forks from extensive degradation. In contrast, efficient replication restart is dependent on Rad51 strand exchange activity, but can be partially rescued by strand exchange-independent mechanisms such as regression of the reversed fork by branch migration or replication origin firing ^52^. Similarly, the rad51-II3A mutant protects stalled replication forks from nucleolytic degradation in *S. pombe* ^53^.

We introduced the *rad51-*II3*A* allele in the strain containing the *ade2* inverted repeats and 14 *Ter* repeats in the blocking orientation. In the absence of Tus, the frequency of spontaneous recombination decreased from 0.62% in the WT to 0.21% in the mutant (Fig 3A, blue data points). Furthermore, induction of fork stalling did not stimulate recombination between the *ade2* inverted repeats (Fig 3A, red data points). We note that although not statistically significant (p-value=0.3), spontaneous and replication-associated recombination in the *rad51Δ* strain was a little higher than in the *rad51-*II3A mutant which might indicate that the presence of inactive rad51-II3A filaments limits recombination events.

**Figure 3.**
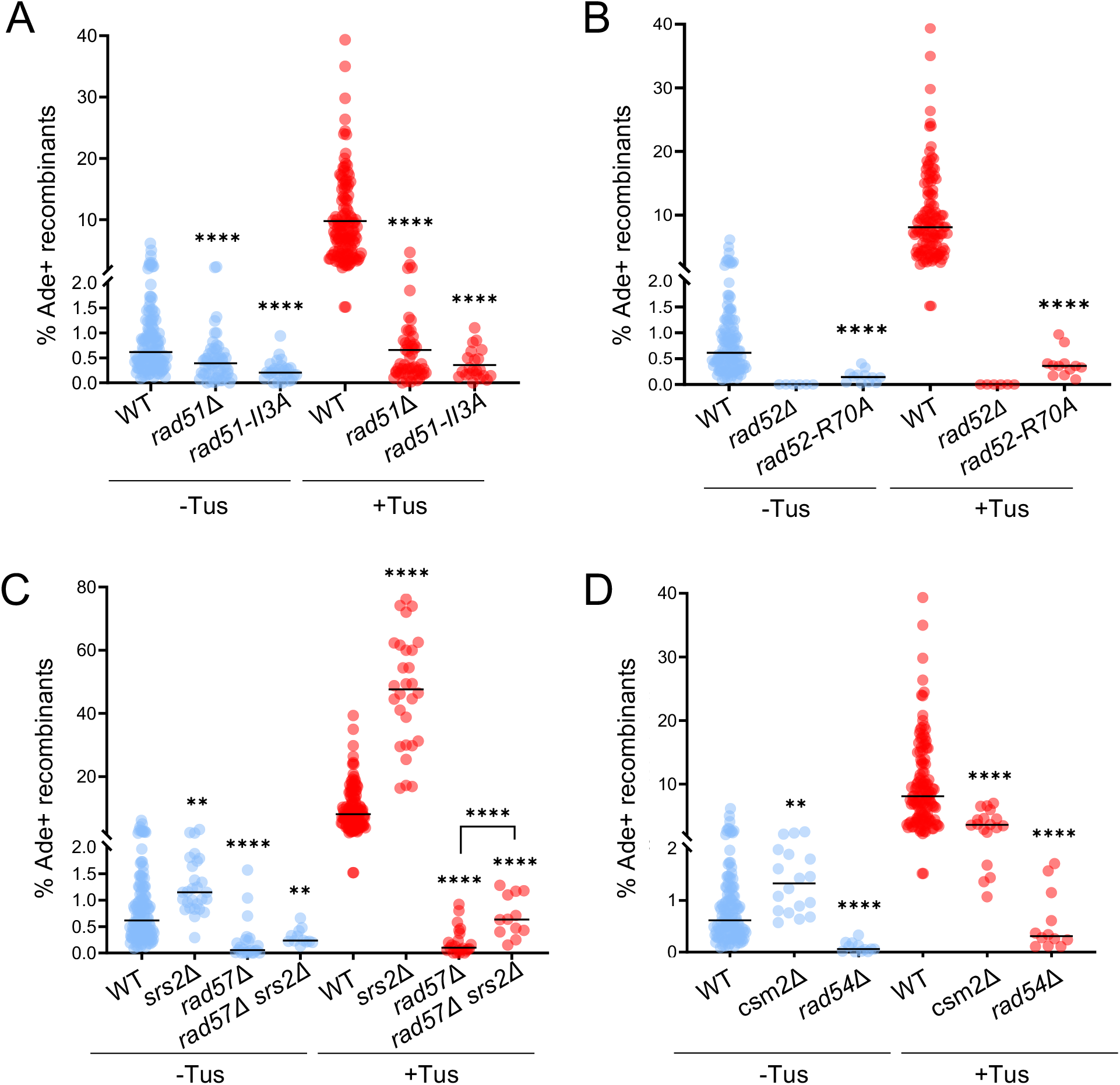
Rad51 strand invasion, Rad52 strand annealing and recombination mediators are required for Tus/*Ter* induced NAHR. **A** Ade^+^ recombination frequencies for WT and *rad51* mutant strains containing 14 *Ter* repeats in the blocking orientation. **B** Ade^+^ recombination frequencies in WT and *rad52* mutant strains containing 14 *Ter* repeats in the blocking orientation. **C** Ade^+^ recombination frequencies in WT, *srs2*Δ and *rad57*Δ single and double mutant strains containing 14 *Ter* repeats in the blocking orientation **D** Ade^+^ recombinants frequencies in WT, *cms2*Δ and *rad54*Δ mutant strains containing 14 *Ter* repeats in the blocking orientation. Black lines indicate medians. P-values, reported as stars when significant, are relative to the WT strain in the same condition: ** p-value<0.005, *** p-value <0.001, **** p-value <0.0001.

Rad52 has two functions in homologous recombination: mediation of Rad51 nucleoprotein filament assembly on RPA-coated ssDNA and annealing of complementary ssDNA during second end capture or SSA at DSBs ^38^. The *rad52-*R70A separation-of-function mutant is proficient for Rad51 loading but defective for ssDNA annealing ^54^ (Fig 3C). We observed a 22-fold decreased frequency of Tus-induced recombination in the *rad52-*R70A mutant strain, consistent with an important role for strand annealing during replication-associated NAHR (Fig 3D).

Taken together, these results suggest that HR associated with fork stalling relies on Rad51-catalyzed strand invasion, distinct from its role in protecting stalled forks from degradation, as well as Rad52-Rad59 catalyzed strand annealing.

### Spontaneous and replication associated NAHR involve different Rad51 mediators

We next assessed the contribution of various Rad51 mediators in spontaneous and Tus/*Ter-*induced recombination. The Rad51 paralogs, Rad55 and Rad57, form a stable heterodimer which assists Rad51 nucleation on RPA-coated ssDNA and promotes rapid re-assembly of filaments after their disruption by the anti-recombinase Srs2 ^25,55,56^. Spontaneous recombination between the *ade2* repeats was reduced 11-fold in the *rad57Δ* mutant strain, consistent with a previous study (Fig 3C, blue data points) ^57^. This finding could indicate that in the absence of the Rad55-Rad57 complex to stabilize Rad51, unstable Rad51 filaments are unable to mediate gene conversion but also inhibit the Rad51-independent spontaneous inversion pathway. When replication fork stalling was induced, there was no stimulation of recombination in the *rad57Δ* strain (Fig 3C, red data points) and recombination was again more deficient in the *rad57Δ* strain than it was in the *rad51Δ* mutant (0.09% vs 0.32%). We also tested whether loss of Srs2 suppresses the *rad57Δ* defects in Tus/*Ter*-induced recombination. Consistent with previous studies ^58^, spontaneous recombination was increased in the *srs2Δ* mutant, and Tus/*Ter* stimulated recombination was increased by 3-fold over the WT value (Fig 3C). Loss of Srs2 partially rescued the *rad57Δ* recombination defect, but the frequency was still 10-fold lower than WT cells, indicating that Rad57’s function is not restricted to antagonizing Srs2.

The Shu complex is another mediator of Rad51 presynaptic filament formation, which interacts directly with Rad51 and the Rad55-Rad57 complex, and has been specifically implicated in the repair of DNA replication-associated damage ^59–63^. Csm2 is one of the four members of the Shu complex. Unlike in the *rad57Δ* mutant, spontaneous recombination between the repeats was not diminished and was even moderately enhanced in the *csm2Δ* mutant (Fig 3D, blue data points). However, when replication fork stalling was induced, recombination in the *csm2Δ* mutant was two-fold lower than in the WT strain (Fig 3D, red data points) suggesting that the Shu complex facilitates NAHR at stalled replication forks but is not strictly required.

Rad54 is an ATP-dependent dsDNA translocase that is required to facilitate Rad51-mediated strand invasion ^64–66^. Consistent with a previous study ^57^, we observed that spontaneous recombination was significantly reduced in the *rad54Δ* mutant, and Tus-induced events were 26-fold lower than WT (0.31% vs 8.08%), similar to the frequency observed for the *rad51Δ* mutant (Fig 3D).

Our results show that Rad51 and its mediators are differentially implicated in spontaneous and replication-associated inverted-repeat recombination. These data indicate that replication-associated NAHR must involve invasion from ssDNA from one *ade2* copy into dsDNA from the other *ade2* copy. Only the long *ade2-n* allele can be restored to a functional *ADE2* gene, so we reasoned that the truncated copy must be the one invaded and used as donor template. Based on the position of the replication fork barrier in our genetic system, fork reversal would promote reannealing of the parental strands of the truncated *ade2* copy, thus providing a dsDNA substrate for invasion.

### Is fork reversal required for replication associated NAHR?

To determine the role of fork reversal in Tus/*Ter*-stimulated recombination we eliminated DNA remodelers that have been implicated in fork reversal. The translocase Rad5 (HLTF in human) initiates replication fork reversal by remodeling the leading strand and proximally positioning the leading and lagging arms, which converts the arrested fork into a chicken-foot structure ^67,68^. However, deletion of *RAD5* showed no significant effect on spontaneous or replication-associated recombination (Fig 4A). The Mph1 helicase (FANCM in human, Fml1 in *S. pombe*) also promotes fork reversal *in vitro* and is required for recombination at a protein-induced fork barrier in *S. pombe* ^69–71^. Loss of Mph1 did not reduce the frequency of spontaneous recombination; however, Tus/*Ter*-stimulated recombination was moderately reduced (p-value=0.05), and recombination was further reduced in the *mph1Δ rad5Δ* double mutant (Fig 4A). Taken together, our results suggest that NAHR events associated with fork stalling require remodeling activity of Mph1 with Rad5 serving a minor or redundant function. Physical analysis of independent recombinants in the *mph1Δ* mutant showed a distribution of replication associated-events similar to the WT strain (Fig S3A). However, in the *mph1Δ rad5Δ* double mutant conversions were reduced 15-fold compared to the WT strain, whereas inversions were only reduced 4-fold (Fig S3A). This could be due to an additional effect of Mph1, in this context, in dissociating the migrating D-loop, thus leading to proportionally more inversions in the *mph1Δ rad5Δ* double mutant.

**Figure 4.**
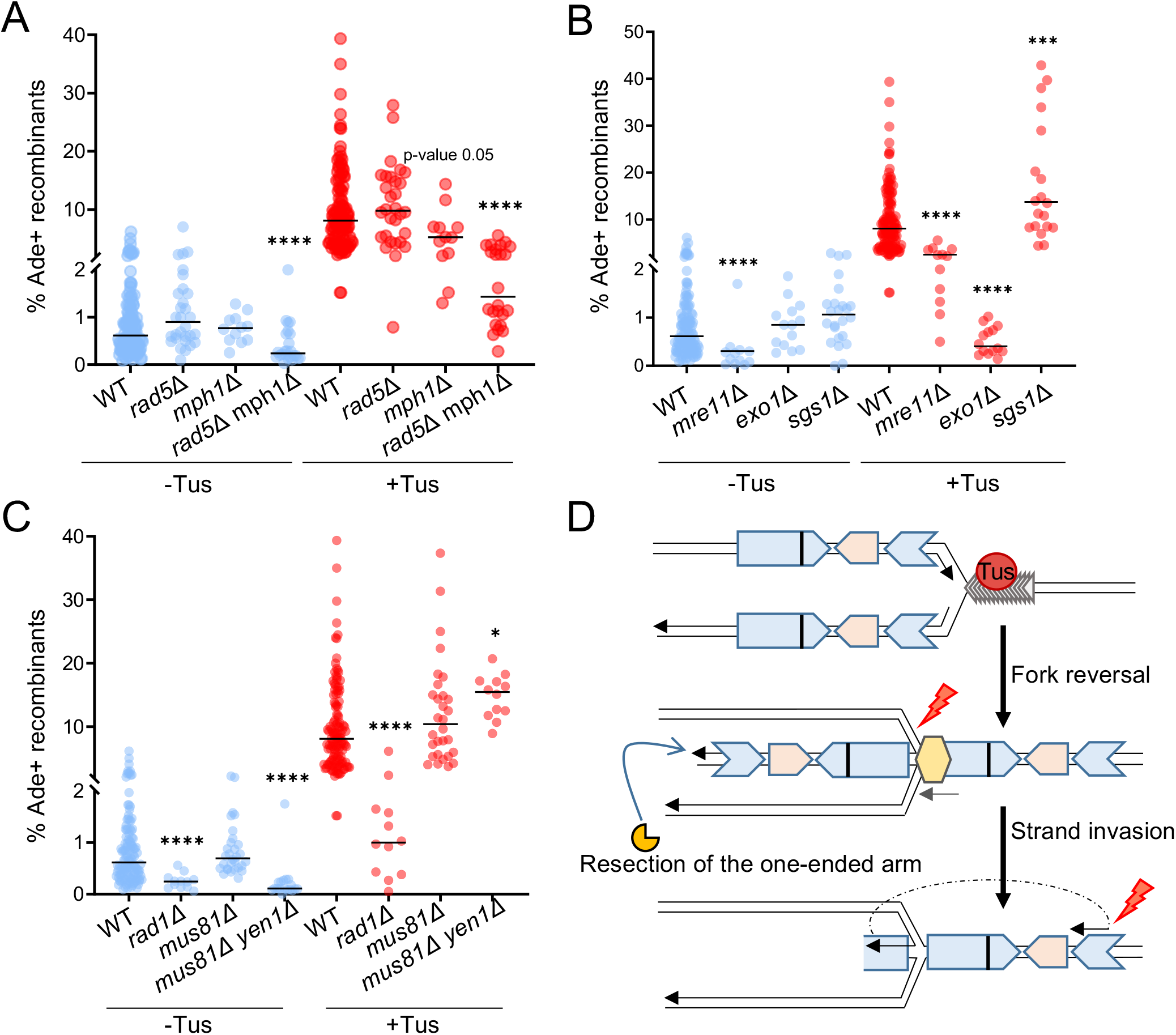
Tus-induced NAHR relies on fork reversal and end resection. **A** Frequency of Ade^+^ recombinants in WT and fork remodeler mutant strains. Black lines indicate medians. P-values, reported as stars when significant, are relative to the WT strain in the same condition: * p-value<0.05, ** p-value<0.005, *** p-value <0.001, **** p-value <0.0001. **B** Ade^+^ recombination frequencies in WT and end resection mutants. **C** Ade^+^ recombination frequencies in WT and nuclease mutants **D** Schematic of stalled forks remodeling. The light yellow hexagon represents the fork reversal mediators Rad5 and/or Mph1. The end resection proteins Mre11 and Exo1 are represented in dark yellow. The red lightning bolt symbolizes cleavage of the recombination intermediates by structure-selective nucleases.

### Tus/Ter stimulated recombination requires Mre11 and Exo1

Fork reversal at the Tus/*Ter* stall, is predicted to form a single-end DSB for end resection to generate a ssDNA substrate for Rad51. DNA end resection occurs by a two-step mechanism involving sequential action by short-range and long-range resection nucleases ^72^. Mre11 nuclease initiates end resection at DSBs as part of the Mre11-Rad50-Xrs2 complex, while Exo1 or Dna2-Sgs1 promotes extensive resection. Studies in *S. pombe* and in mammalian cells have shown that the same nucleases can degrade regressed forks ^4,73^. In budding yeast, MRX is essential for resection of DSBs with end-blocking lesions, but resection can still occur at “clean” DSBs by the direct action of the long-range nucleases, Exo1 and Dna2 ^72^.

To determine the role of DNA end resection in Tus/*Ter*-stimulated recombination, we eliminated DNA nucleases that function in short-range and long-range resection. The frequency of spontaneous and Tus-induced recombination was reduced by 3-fold in the *mre11Δ* mutant (Fig 4B) indicating a role for resection initiation by MRX. In the absence of Exo1, spontaneous recombination occurred at the WT level (Fig 4B, blue data points); however, stimulation of recombination by the Tus/*Ter* barrier was abolished (Fig 4B, red data points). This finding suggests that replication-associated NAHR relies on extensive degradation of the newly synthesized lagging strand by Exo1 to generate a ssDNA leading strand substrate for Rad51 loading.

Sgs1 and Dna2 act redundantly with Exo1 in long-range resection at DSBs. The striking defect in Tus/*Ter*-stimulated recombination in the *exo1Δ* mutant suggests that Exo1 plays a more important role in fork resection than Sgs1-Dna2. Consistent with this interpretation, we found that the frequency of Tus/*Ter*-stimulated recombination was not reduced in the *sgs1Δ* mutant compared to the WT (Fig 4B, red data points). In the absence or presence of Tus expression, *sgs1Δ* cells showed a slight increase in recombination (Fig 4B, blue data points), consistent with the previously reported hyper-recombination phenotype ^74^. This result seems to indicate that Sgs1 does not play a significant role in fork resection. However, the caveat is that Sgs1 is involved in other processes, such as the dissolution of recombination intermediates, and these roles could mask a role in fork resection ^75^.

### Opposing roles of structure-selective nucleases in replication associated NAHR

Fork reversal at the Tus-induced barrier could generate an invading end with a short sequence heterology that would need to be removed to prime DNA synthesis within the D-loop intermediate. Previous studies in yeast have shown that Rad1-Rad10 nuclease removes 3′ heterologies during Rad51-dependent strand invasion, as well as 3′ flaps formed during Rad51-independent SSA ^76^. Consistent with the need for heterologous flap or loop removal, the frequency of Tus/*Ter*-induced recombination was significantly reduced in the *rad1Δ* mutant (Fig 4C, red data points).

Fork reversal creates a four-way junction that can be cleaved by structure-selective nucleases to create a one-ended DSB. In budding yeast, Mus81-Mms4 is the main nuclease responsible for cleaving recombination intermediates, with Yen1 providing a back-up function ^5,77–79^. We did not find a significant change in the frequency of spontaneous or replication-associated recombination in the *mus81Δ* mutant. However, elimination of Yen1 and Mus81 resulted in a 2-fold increase in the frequency of Tus/*Ter-*induced recombination from 8.08% to 15.45% (Fig 4C). Thus, Mus81-Mms4 and Yen1 may abort the normal process for forming recombinants at the Tus/*Ter* barrier (Fig 4D). We also looked at the distribution of replication associated-recombination events in the *mus81Δ yen1Δ* double mutant (Fig S3B). Inversions represent more that 50% of the products in the double mutant indicating that they are not generated by cleavage of a HJ-containing intermediate. The increase in Tus/*Ter*-stimulated inversion products in the *mus81Δ yen1Δ* double mutant suggests that Mus81-Mms4 and Yen1 might cleave the migrating D-loop initiated by Rad51, in addition to the reversed fork intermediate (Fig 4D).

### A specific role for the replicative polymerase Pol δ at replication associated NAHR

NAHR is predicted to require DNA synthesis to convert the *ade2-n* allele, and potentially to invert the *TRP1* gene between the repeats. *In vivo* and *in vitro* studies have shown that DNA Pol δ initiates synthesis from the invading 3’ end within the D-loop intermediate ^80–83^. *S. cerevisiae* Pol δ is a heterotrimer comprised of a catalytic subunit Pol3 and two accessory subunits Pol31 and Pol32 ^84^. Pol31 and Pol32 also associate with Rev3 and Rev7 to form another B-family DNA polymerase, Pol ζ, a translesion polymerase responsible for mutagenic replication of damaged DNA ^85,86^.

When we deleted *POL32* in the *ade2* reporter strain containing 14 *Ter* repeats in the blocking orientation, we observed a decrease in the frequency of spontaneous recombination (Fig 5, blue data points). Upon induction of fork stalling by Tus/*Ter* in the *pol32Δ* mutant, we observed a significant decrease of recombination compared to the WT strain (2.08% vs 8.08 % in WT) (Fig 5, red data points). To determine whether the *pol32Δ* defect was due to Pol δ or Pol ζ, we measured recombination frequencies in a *rev3Δ* mutant. Unlike the *pol32Δ* strain, the *rev3Δ* mutant showed a full stimulation of recombination upon induction of the Tus/*Ter* barrier. The double mutant exhibited a similar phenotype to the *pol32Δ* single mutant; thus, Pol δ but not Pol ζ appears to be involved in this process.

**Figure 5:**
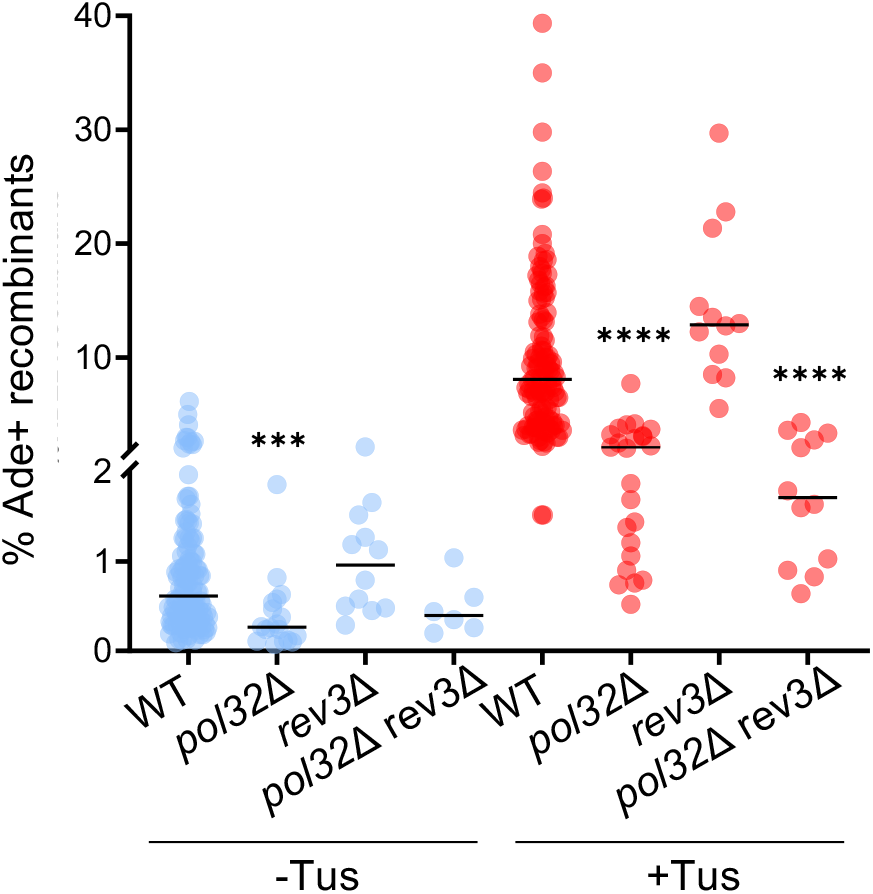
Pol δ catalyzes DNA synthesis during replication associated-NAHR. Ade^+^ recombination frequencies in WT and DNA polymerase mutants. Black lines indicate medians. P-values, reported as stars when significant, are relative to the WT strain in the same condition: *** p-value <0.001, **** p-value <0.0001.

## DISCUSSION

Replication stress, defined as a slowing down or complete arrest of DNA synthesis during chromosome replication, has emerged as a primary cause of genome instability, a hallmark of cancer and other human disorders associated with genomic rearrangements ^1,87,88^. In fission yeast and mammalian cells, replication fork stalling adjacent to a recombination reporter can lead to increased recombination events ^44,89,90^. In this work, we show that a Tus/*Ter* barrier designed to induce transient replication fork stalling near inverted repeats stimulates recombination mediated by a unique genetic pathway, distinct from spontaneous NAHR or post-replicative repair. The model presented in Figure 6, which is discussed in detail below, is based on our genetic findings and builds on other template switching models for replication-associated recombination.

**Figure 6:**
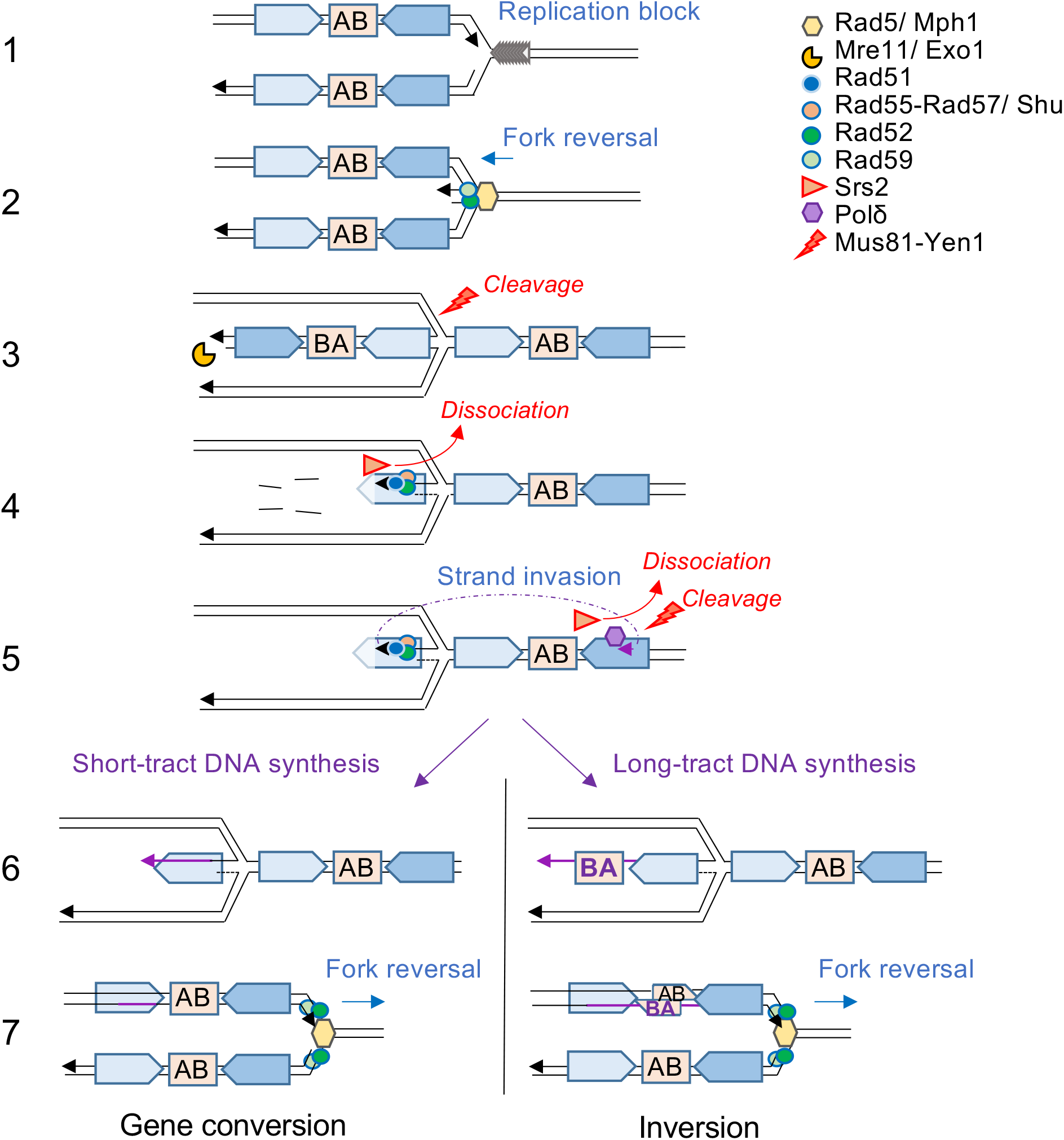
Model for NAHR at arrested replication forks. 1) The replication fork stalls at the Tus/*Ter* barrier. The letters AB symbolize the orientation of the intervening sequence. 2) Rad5 and Mph1 catalyse fork reversal. Rad52-Rad59 annealing activity could facilitate strand pairing of the daughter or parental strands 3) The regressed arm is degraded by Mre11 and Exo1 nucleases. The reversed fork can be cleaved by nucleases, aborting NAHR. 4) Rad51 polymerization on ssDNA is mediated by Rad52 and Rad55-Rad57 (with help from the Shu complex), counteracting the Srs2 anti-recombinase. 5) Rad51 catalyses strand invasion into the parental non-allelic repeat, heterologies are cleaved by Rad1-Rad10 and DNA synthesis is initiated by Pol δ. The D-loop can be dissociated by Mph1 or Srs2, or cleaved by Mus81-Mms4 and Yen1. 6) Short tract DNA synthesis leads to gene conversion, whereas long tract DNA synthesis leads to inversion of the intervening sequence (AB -> BA) on the newly synthesized leading strand. 7) Regression of the reversed fork is necessary to restart replication. The large unpaired heterology loop could be repaired by Rad1-Rad10 or segregated at the next replication cycle.

### Replication-associated NAHR has different genetic requirements to spontaneous NAHR and post replicative repair pathways

Spontaneous inversion of the sequence between inverted repeats is dependent on Rad52 and Rad59, whereas spontaneous gene conversion without an associated inversion is dependent on Rad52 and Rad51 ^39,42^. In contrast, we show here that HR induced at a replication-fork barrier triggers both inversions and gene conversions mediated by Rad51, Rad59 and Rad52 working together in a unique pathway.

The template switching mechanism of post replicative repair (PRR) is a DNA damage tolerance pathway that involves use of the undamaged sister chromatid as a homologous template for lesion bypass ^7,91^. One model of template switching involves reversal of the stalled fork for stabilization and/or repositioning of the lesion to bypass damage ^9,10,92^. The other model involves pairing of a template strand at a ssDNA gap with the undamaged sister chromatid to form a pseudo-double Holliday junction (dHJ) intermediate ^7^. The second mode of PRR template switching is mediated by several proteins that we show are also important for NAHR at Tus/*Ter* stalled forks: Rad51, Rad55-Rad57, Csm2, Exo1 and DNA Pol δ ^93–95^. However, Rad59 is not required for PRR template switching, whereas we detected a strong reduction of NAHR at stalled forks in the *rad59Δ* mutant ^93^. In addition, Rad5, which is essential for PRR template switching ^94^, is only required for Tus-induced recombination in the absence of Mph1 (see later paragraph). Overall, our data show that the mode of HR associated with replication stalling at repetitive sequences is genetically different from spontaneous HR or PRR template switching pathways.

### Why are Rad59, Rad52 and Rad1 required for Rad51-mediated NAHR at stalled forks?

Rad59 contributes to a subset of HR events by assisting Rad52 in second end capture during DSB repair and in SSA at direct repeats ^96,97^. Thus, Rad59-dependent recombination is thought to be linked to DSB repair where both ends have to be rescued through simultaneous interactions with an unbroken template. However, reversal of the stalled fork at the Tus/*Ter* barrier would generate a regressed arm resembling a one-ended break with no second end for capture by Rad52-Rad59 mediated strand annealing. Thus, the sizable decrease in Rad51-mediated HR at Tus/*Ter* in the *rad59Δ* mutant is intriguing. We suggest that Rad59 acts with Rad52 in facilitating regression of the stalled fork, or restoration of the reversed fork, by mediating annealing of nascent or parental strands. Interestingly, the role of mammalian RAD52 has long remained mysterious due to the presence of BRCA2, which assumes RAD51-mediator function and prevents any significant DNA repair phenotype in RAD52-deficient cells. But RAD52 was recently shown to have a specific protective role in maintaining cell viability under replication stress that is non-redundant with BRCA2 ^98,99^. Our results suggest a conserved function for Rad52 during replication stress, involving its strand annealing activity.

Rad59 could also function by stabilizing an annealed intermediate with a heterologous tail for cleavage by Rad1-Rad10, as previously suggested ^100^. Such an intermediate could occur after fork resetting (Fig 6). The other possible functions of Rad1-Rad10 could be in repair of the large loop heterology expected to occur from long tract synthesis and fork reset ^101^; however, we would not expect loss of this function to reduce the frequency of Ade^+^ recombinants.

### Rad5 and Mph1 redundantly mediate fork reversal at Tus/*Ter* stalled replication forks

We did not detect a decrease of replication associated-NAHR in the *rad5Δ* single mutant, again highlighting the specific genetic requirements of this pathway compared to PRR template switching. However, we found a 5.5-fold decrease in Tus-induced recombination in the *rad5Δ mph1Δ* strain compared to the WT. The relationship between Rad5 and Mph1, the two major DNA remodelers in budding yeast with reported replication fork regression activity, is not fully understood. The additive effect in genotoxic sensitivity observed in the double mutant ^102^, and partial suppression of MMS sensitivity of a *rad5Δ* mutant by Mph1 hyperactivation, suggests that they have overlapping activities ^103^, consistent with our findings. The requirement for Rad51 strand invasion activity leads us to propose a model involving invasion of the parental duplex, which would require fork regression to create an invading end. We note that the fork would need to reverse by several kb for the *ade2-n* allele to be placed for invasion of the reformed parental *ade2-5′Δ* allele (Fig 6), which could lead to an under-estimation of recombination induced by the Tus/*Ter* block. Mph1 is also involved in D-loop dissociation during DSB repair and HR-mediated restart of collapsed replication forks ^104–106^. If the main activity of Mph1 during NAHR at stalled forks is to dissociate the D-loop we would not expect to observe a reduction in recombination frequency, although the change in the proportion of inversions in the *rad5Δ mph1Δ* double mutant is consistent with D-loop dissociation activity of Mph1. It was recently proposed that Mph1 can act coordinately with Rad54 and Rad5 in the HR-driven fork regression mechanism to bypass stalled replication forks ^107^. We observed a strong reduction of spontaneous and replication-associated NAHR events in the *rad54Δ* mutant. This outcome could be due to a role in fork reversal in addition to the role of Rad54 in promoting Rad51-mediated strand invasion ^64^. However, based on the phenotype of *rad5Δ mph1Δ* double mutant, Rad54 does not appear to play a major role in fork reversal.

### A model for inverted repeat recombination at stalled forks

We envision that a stalled replication fork is reversed into a chicken foot structure by the redundant activities of Rad5 and Mph1 (and potentially Rad54). We propose that the process is assisted by Rad52 and Rad59 which facilitate nascent strand pairing. Fork reversal creates a branched structure that could be acted upon by endonucleases such as Mus81-Mms4 and Yen1 or counteracted by helicases. The reversed fork exposes a regressed arm which is processed to form a 3’ ssDNA overhang by the sequential activities of Mre11 and Exo1, generating a ssDNA template for Rad51 nucleoprotein filament formation on the leading strand. The resection activity of Exo1 is redundant with that of Sgs-Dna2 during DSB repair but we did not observe a defect in NAHR events at stalled forks in the *sgs1Δ* mutant. Another activity of Exo1 shown in human, is to recruit translesion synthesis DNA polymerases to sites of damage ^108^. However, our results indicate that the translesion polymerase Pol ζ is not involved in DNA synthesis during NAHR at stalled forks.

We show that Rad51 loading on the leading ssDNA template is facilitated by Rad55-Rad57 and the Shu complex. One activity of Rad55-Rad57 is to counteract the anti-recombinase Srs2, but our finding that deletion of *SRS2* only partially suppresses the *rad57Δ* HR defect suggests an additional function for the Rad51 paralog complex. Interestingly, a recent study showed that Rad55-Rad57 is essential for the promotion of UV-induced HR independently of Srs2 and prevents the recruitment of translesion synthesis polymerases which would compete with template switching ^109^.

We propose that Rad51 catalyzes strand invasion into the parental non-allelic inverted sequence ahead of the reversed fork, facilitated by the dsDNA translocase Rad54. DNA synthesis is mediated by Pol δ using the repetitive sequence as a template. Dissociation of the extended invading strand prior to the intervening sequence (represented by AB in Fig 6) would result in no inversion. Such dissociation may be promoted by Mph1. On the other hand, long tract DNA synthesis through the intervening sequence (B before A on Fig 6) could result in its inversion. Theoretically, inversions could also result from cleavage of a Holliday junction intermediate by Mus81-Mms4 or Yen1. However, we did not observe any decrease of inversions in the *mus81Δ yen1Δ* double mutant (Fig S3B).

The reversed fork would then need to be regressed by the action of remodelers and/or strand annealing proteins to restore the replication fork. Regression of the reversed fork could dissociate the D-loop, or helicases could dismantle the D-loop prior to regression. The resulting replication fork would contain heteroduplex DNA encompassing the *ade2-n* allele with the potential to create a functional *ADE2* gene by mismatch correction or segregation of the strands at the next cell cycle. A heterologous tail, or loop, formed between the inverted repeats by long tract synthesis could be cleaved by Rad1-Rad10, or segregate at the next replication cycle resulting in two daughter cells with either a conversion or an inversion.

In conclusion, this work uncovers a genetically unique pathway that is stimulated by localized replication stress and can mediate genomic rearrangements of repetitive sequences. It should be noted that our reporter system can only reveal recombination events that lead to the restoration of a functional *ADE2* gene. Reactions where strand invasion of the non-allelic copy occurred downstream from the +2 frameshift location would not be detected. The frequency of NAHR events at inverted repeats that can generate rearrangements is thus likely to be underestimated in our assay.

How spontaneous Rad51-independent inversions are generated remains an open question to be explored. Previous studies suggest they are not due to DSB repair and this work supports the idea they are not associated with fork stalling at a protein barrier. Other contexts that could be investigated are replication uncoupling to form long stretches of ssDNA and fork collision with the transcription machinery.

## METHODS

### Yeast strains

All yeast strains are derived from W303, corrected for the *rad5-535* mutation, and are *ade2::hisG* (Table S2). The *ade2* inverted-repeat recombination reporter was described previously ^39^. In this study, the reporter was amplified by 2-rounds PCR from the strain 2002-9D ^42^ and integrated at the *his2* locus on chromosome 6.

The 14 *TerB* repeats were amplified by PCR from plasmids pNBL63 (blocking orientation) and pNBL55 (permissive orientation) and integrated 170 bp and 120 bp distal to the *ade2Δ5’* repeat, respectively. The *PGAL1-*HA-*Tus* cassette was cloned from plasmid p415-*PGAL1-*HA-*Tus* ^43^ into pRG205MX ^110^, adjacent to the yeast *LEU2MX* selectable marker, and integrated at the *LEU2* locus.

All mutant strains were constructed by genetic crosses using haploid strains in the laboratory collection, with the exception of *csm2Δ*, *mph1Δ* and *mus81Δ* strains that were obtained by transformation with a *KanMX* ORF replacement cassette. Strains used for 2D gels additionally contained a *bar1::HphMX* allele generated by transformation or by genetic cross.

### Measurement of recombination frequency

The percentage of Ade^+^ recombinants, which corresponds to recombination frequency, was measured as follows. Strains were grown for 3 days on YPAD (1% yeast extract, 2% bacto-peptone, 2% dextrose, 10mg/L adenine) or 4 days on YPAG (1% yeast extract, 2% bacto-peptone, 2% galactose, 10mg/L adenine) plates. Colonies of similar size (described below as initial colonies) were suspended in 1mL water to an OD_600_ close to 0.3. Cells were serially diluted and plated on YPAD or synthetic complete – adenine (SC-Ade) medium. Colonies were counted 2 days after plating and two dilutions from each initial colony were averaged. The percent Ade^+^ recombinants was determined by the ratio of the number of colonies growing on SC-Ade plates and YPAD plates x 100. Each data point in the graphs shows the percentage of Ade^+^ recombinants measured from one initial colony. The medians, shown on graphs as black lines, were calculated for each strain and condition from multiple independent trials and are indicated in Table S1, as well as the number of initial colonies tested.

### Distribution of Ade^+^ recombinants

To ensure analysis of independent NAHR events, only one Ade^+^ recombinant colony from each initial colony (see above) was used. Inversions and conversions were scored by PCR using a primer annealing to the *his2* sequence upstream of the *ade2* reporter or to the *ade2-n* cassette, and primers of opposite orientation that anneal to the *TRP1* sequence between the repeats (see fig 1E). The number of independent recombinants tested for each strain and condition is indicated in the figures.

### Statistical analysis

Ade^+^ recombination frequencies were analyzed on log transformed values by one-way Anova with a Boneferroni post-test. Spontaneous and Tus/*Ter* associated data were analyzed separately. Distributions of inversions and conversions among Ade^+^ recombinants were analyzed by Chi-Square test. Stars indicate a significant difference with the WT strain in the same condition: * p-value <0.05, ** p-value <0.005, *** p-value <0.001, **** p-value <0.0001. Where relevant, exact p-values are indicated on figures.

### Two-dimensional (2D) gel analysis of replication intermediates

Yeast cultures were grown overnight in YEPL (1% yeast extract, 2% bacto-peptone, 10 mg/L adenine, 3% sodium DL-lactate) medium to OD_600_= 0.8. Cultures were synchronized in G1 with 1.5 μg/ml alpha factor mating pheromone (GenScript) for 3 h at 30°C. Tus expression was induced by adding 2% Galactose (final w/v) for the final 2.5 h of the G1-arrest. Cells were released from G1-arrest by centrifugation, washing and resuspension in warm YEPL medium containing 100 μg/mL pronase. Arrest and release of the cultures were checked by flow cytometry. Cells were incubated for 50 minutes at 30°C, then cultures were placed on ice and treated with 0.1% sodium azide to stop metabolism. The hexadecyltrimethylammonium bromide (CTAB) protocol was followed for extraction of total genomic DNA ^111^. A Qubit Flex fluorometer (Invitrogen) was used for quantification and the DNA yield was about 30 μg from each 200 mL overnight culture.

For each 2D gel, 15 μg of genomic DNA was digested overnight with 90 U ClaI. Samples were run on the first-dimension gel (0.35% agarose, 1x Tris-Borate-EDTA) at constant voltage of 1V/cm for ~ 19h, and then stained with 0.3 μg/mL ethidium bromide. Gel strips were excised under a UV trans-illuminator, rotated by 90° and run on a second gel (1.15% agarose, 1x Tris-Borate-EDTA, 0.3 μg/mL ethidium bromide) at 4V/cm for ~ 6h at 4°C.

### Southern blotting

After denaturation and neutralization of the gels, DNA was transferred in 2 x SSC to positively charged nylon membranes (GE Healthcare Amersham Hybond-N+) and was then immobilized by ultraviolet cross-linking (1200 J). DNA fragments were detected using a mix of five probes labelled by PCR amplification with ^32^P dCTP (Perkin Elmer) described in Fig S1. ULTRA-hyb Ultrasensitive hybridization buffer (Invitrogen) was used for hybridization of the probes at 42°C. Membranes were washed as recommended by the manufacturer. 2D gels were exposed for 4 hours in a phosphor screen cassette and the signal was detected with a Typhoon Trio phosphoimager (GE healthcare).

## ACKNOWLEDGEMENTS

We thank H. Mankouri and R. Rothstein for generous gifts of plasmids and strains, and W.K. Holloman, R. Reid and members of the Symington lab for comments on the manuscript. This work was supported by grants from the National Institutes of Health (R21 ES030447 and R35 GM126997 to L.S.S.)

## AUTHOR CONTRIBUTIONS

L.M. performed all experiments. L.M. and L.S.S. contributed to the study design, data analysis and manuscript preparation.

---

**Figure S1 :**
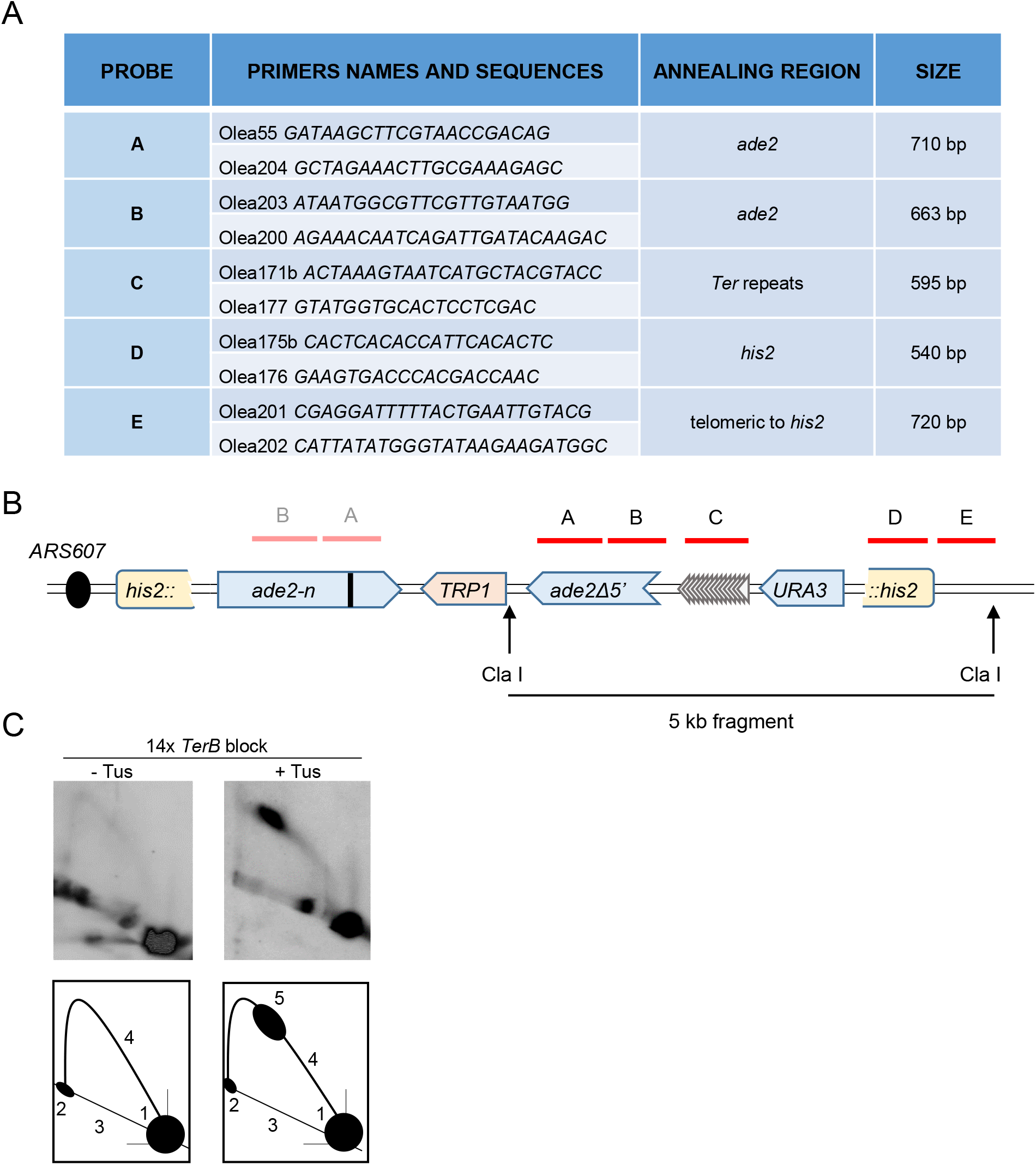
Two-dimensional gel analysis of replication intermediates. **A** Table of oligonucleotides used to generate radiolabelled probes for hybridization. **B** Schematic of the digestion fragment detected by 2D gel. Radiolabelled probes are represented by red lines and cover 3.2 kb of the digested 5 kb fragment. Probes A and B anneal to the two *ade2* repeats. After Cla I digestion, the *ade2-n* repeat is part of a 11.2 kb fragment which does not interfere with detection of the signal from the smaller 5kb fragment containing the *Ter* repeats. **C** Interpretation of 2D gel analysis images. 1 = Cla I fragment before replication. 2 = Cla I fragment after full replication. 3 = non specific linear DNA. 4 = arc of Y-shaped replication intermediates. 5 = fork stall corresponding to the position of the Tus/*Ter* barrier.

**Figure S2:**
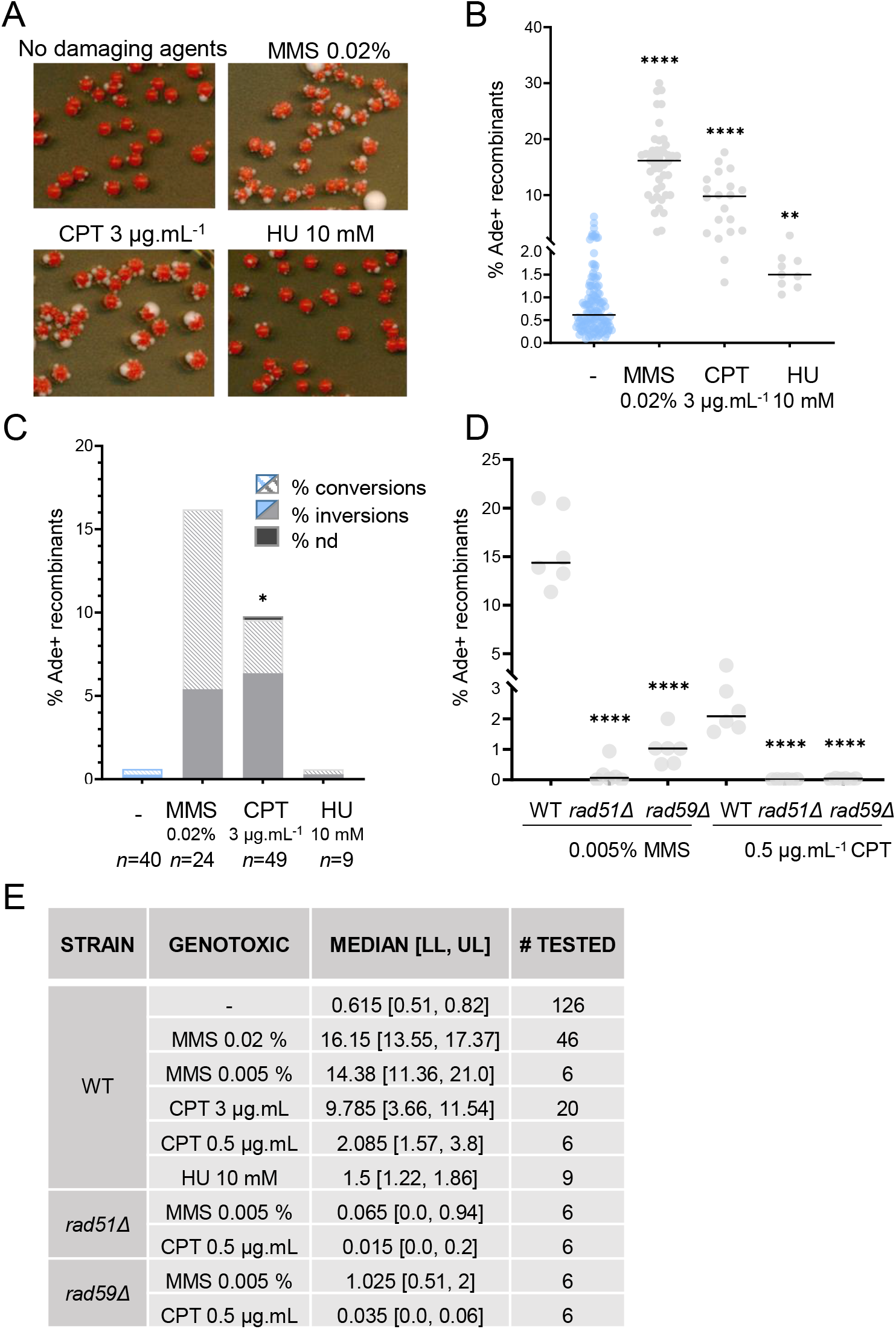
Genome wide replication stress induces Rad51 and Rad59-dependent NAHR at inverted repeats. **A** Colonies form more white sectors (indicative of Ade ^+^ phenotype) on plates containing genotoxic agents inducing replication stress. **B** Ade^+^ recombination frequencies without (blue data points) and with (grey data points) genotoxic agents. Concentrations from 2 mM to 150 mM HU were tested and showed comparable results. Black lines indicate medians. P-values, reported as stars when significant, are relative to the NO genotoxic agents data: **** p-value <0.0001, ** p-value<0.005. **C** Distribution of NAHR events, with and without genotoxic agents, scored by PCR. Independent events were examined for each strain. Striped color = conversions, plain color = inversions, nd = not determined by PCR. P-values, reported as stars when significant, are relative to the NO genotoxic agents data: * p-value <0.05 **D** Ade + recombination frequencies with low concentrations of MMS and CPT in the WT and mutant strains. Black lines indicate medians. P-values, reported as stars when significant, are relative to the WT strain in the same condition: **** p-value <0.0001. **E** Quantification of Ade + recombinants in the different strains in presence of genotoxic agents. >95% confidence intervals to the median are indicated as [LL, UL], where LL is the lower limit and UL is the upper limit.

**Figure S3:**
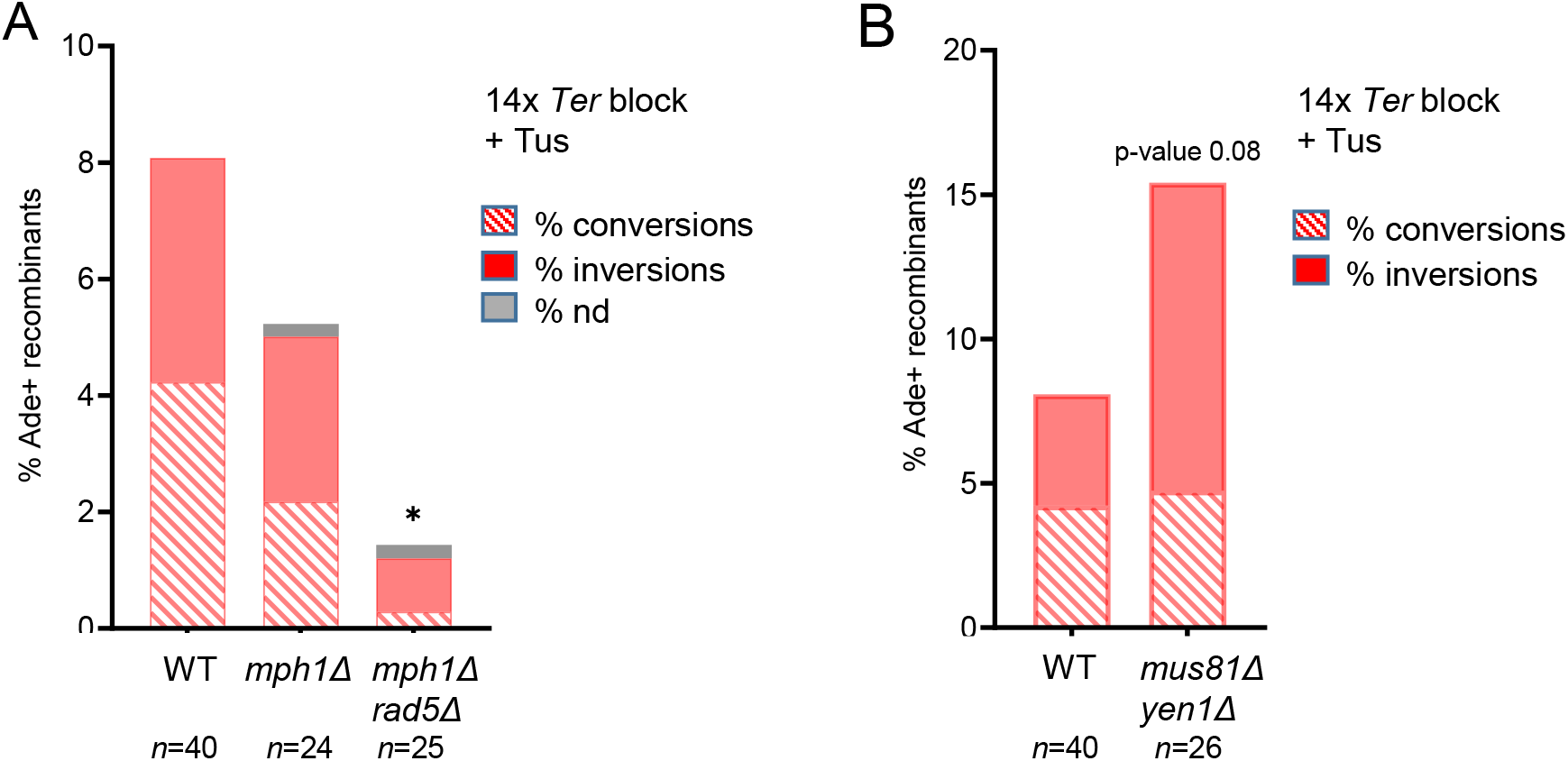
Distribution of NAHR events in WT and mutant strains. **A** Distribution of NAHR events in the WT, *mph1Δ* and *mph1Δ rad5Δ* strains, scored by PCR. **B** Distribution of NAHR events in the WT and *mus81Δ yen1Δ strain*, scored by PCR. nd= not determined by PCR, *n* indicates the number of independent Ade^+^ recombinants tested. P-values, reported as stars when significant, are relative to the WT strain: * p-value <0.05.

**Table S1:**
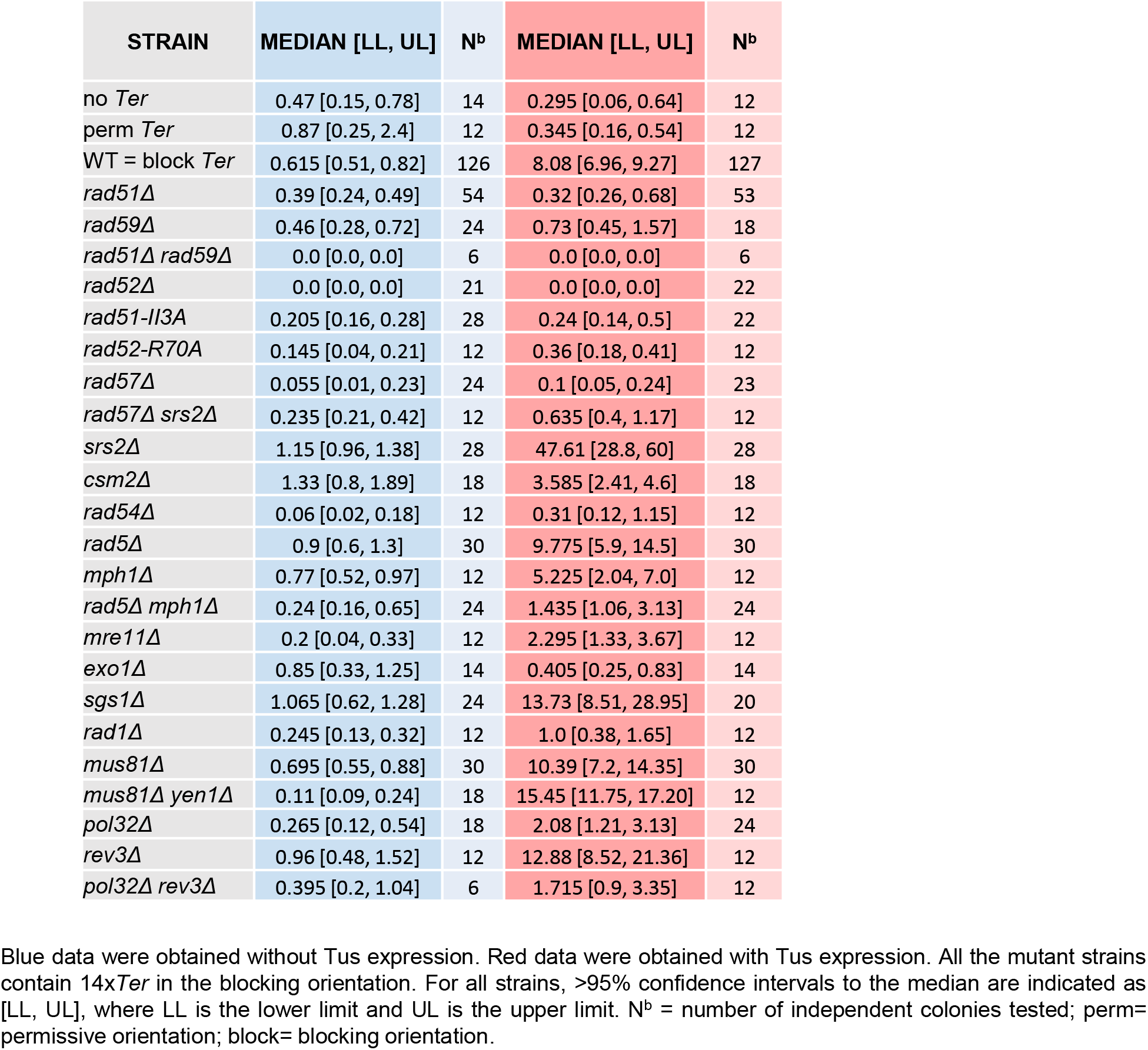
Ade^+^ recombination quantifications.

**Table S2:** Yeast strains.

| STRAIN | RELEVANT GENOTYPE | USE |
| --- | --- | --- |
| LSY2002-9D | *his3::ade2Δ5'-TRP1-ade2-n* | Strain construction |
| LS4040 | *his2::ade2-n-TRP1- ade2Δ5'* | Strain construction |
| LSY4577 | *his2::ade2-n-TRP1- ade2Δ5' leu2::Gal-TUS-LEU2MX* | Recombination assay |
| LSY4579-1 | *his2::ade2-n-TRP1- ade2Δ5'-perm14xTerB-URA3 leu2::Gal-TUS-LEU2MX* | Recombination assay |
| LSY4581-2 | *his2::ade2-n-TRP1- ade2Δ5'-block14xTerB-URA3 leu2::Gal-TUS-LEU2MX* | Recombination assay |
| LSY4583 | *his2::ade2-n-TRP1- ade2Δ5'-block14xTerB-URA3 leu2::Gal-TUS-LEU2MX rad59::LEU2* | Recombination assay |
| LSY4584 | *his2::ade2-n-TRP1- ade2Δ5'-block14xTerB-URA3 leu2::Gal-TUS-LEU2MX rad59::LEU2 rad51::HIS3* | Recombination assay |
| LSY4585 | *his2::ade2-n-TRP1- ade2Δ5'-block14xTerB-URA3 leu2::Gal-TUS-LEU2MX rad51::HIS3* | Recombination assay |
| LSY5127 | *his2::ade2-n-TRP1- ade2Δ5'-block14xTerB-URA3 leu2::Gal-TUS-LEU2MX rad52::TRP1* | Recombination assay |
| LSY4618-1 | *his2::ade2-n-TRP1- ade2Δ5'-block14xTerB-URA3 leu2::Gal-TUS-LEU2MX rad51-II3A-KanMX* | Recombination assay |
| LSY5122 | *his2::ade2-n-TRP1- ade2Δ5'-block14xTerB-URA3 leu2::Gal-TUS-LEU2MX rad57::LEU2* | Recombination assay |
| LSY5134 | *his2::ade2-n-TRP1- ade2Δ5'-block14xTerB-URA3 leu2::Gal-TUS-LEU2MX srs2::TRP1* | Recombination assay |
| LSY5013-12C | *his2::ade2-n-TRP1- ade2Δ5'-block14xTerB-URA3 leu2::Gal-TUS-LEU2MX srs2::TRP1 rad57::LEU2* | Recombination assay |
| LSY5123 | *his2::ade2-n-TRP1- ade2Δ5'-block14xTerB-URA3 leu2::Gal-TUS-LEU2MX csm2::KanMX* | Recombination assay |
| LSY5128-1 | *his2::ade2-n-TRP1- ade2Δ5'-block14xTerB-URA3 leu2::Gal-TUS-LEU2MX rad54::LEU2* | Recombination assay |
| LSY4459-1 | *his2::ade2-n-TRP1- ade2Δ5'-block14xTerB-URA3 leu2::Gal-TUS-LEU2MX rad5::URA3* | Recombination assay |
| LSY5130 | *his2::ade2-n-TRP1- ade2Δ5'-block14xTerB-URA3 leu2::Gal-TUS-LEU2MX mph1::KanMX* | Recombination assay |
| LSY5132 | *his2::ade2-n-TRP1- ade2Δ5'-block14xTerB-URA3 leu2::Gal-TUS-LEU2 mph1::KanMX rad5::URA3* | Recombination assay |
| LSY5124-1 | *his2::ade2-n-TRP1- ade2Δ5'-block14xTerB-URA3 leu2::Gal-TUS-LEU2MX mre11::LEU2* | Recombination assay |
| LSY4620 | *his2::ade2-n-TRP1- ade2Δ5'-block14xTerB-URA3 leu2::Gal-TUS-LEU2MX sgs1::HIS3* | Recombination assay |
| LSY4621-1 | *his2::ade2-n-TRP1- ade2Δ5'-block14xTerB-URA3 leu2::Gal-TUS-LEU2MX exo1::KanMX* | Recombination assay |
| LSY5135 | *his2::ade2-n-TRP1- ade2Δ5'-block14xTerB-URA3 leu2::Gal-TUS-LEU2MX rad1::LEU2* | Recombination assay |
| LSY5129 | *his2::ade2-n-TRP1- ade2Δ5'-block14xTerB-URA3 leu2::Gal-TUS-LEU2MX mus81::KanMX* | Recombination assay |
| LSY5133-1 | *his2::ade2-n-TRP1- ade2Δ5'-block14xTerB-URA3 leu2::Gal-TUS-LEU2MX mus81::KanMX yen1::HIS3* | Recombination assay |
| LSY4592-1 | *his2::ade2-n-TRP1- ade2Δ5'-block14xTerB-URA3 leu2::Gal-TUS-LEU2MX pol32::KanMX* | Recombination assay |
| LSY5125-1 | *his2::ade2-n-TRP1- ade2Δ5'-block14xTerB-URA3 leu2::Gal-TUS-LEU2MX pol32::KanMX rev3::HIS3* | Recombination assay |
| LSY5126 | *his2::ade2-n-TRP1- ade2Δ5'-block14xTerB-URA3 leu2::Gal-TUS-LEU2MX rev3::HIS3* | Recombination assay |
| LSY5067-15 | *his2::ade2-n-TRP1- ade2Δ5'-block14xTerB-URA3 bar1::HYG* | 2D gel analysis |
| LSY5060-4 | *his2::ade2-n-TRP1- ade2Δ5'-block14xTerB-URA3 leu2::Gal-TUS-LEU2MX bar1::HYG* | 2D gel analysis |
| LSY5075 | *his2::ade2-n-TRP1- ade2Δ5'-block14xTerB-URA3 leu2::Gal-TUS-LEU2MX rad51::HIS3 bar1::HYG* | 2D gel analysis |
| LSY5062 | *his2::ade2-n-TRP1- ade2Δ5'-block14xTerB-URA3 leu2::Gal-TUS-LEU2MX rad59::LEU2 bar1::HYG* | 2D gel analysis |

All strains are haploids derived from W303 (*leu2-3,112 trp1-1 can1-100 ura3-1 his3-11,15*), *ade2::hisG* and *RAD5* unless otherwise indicated.

